# Isotope Encoded chemical Imaging Identifies Amyloid Plaque Age Dependent Structural Maturation, Synaptic Loss, and Increased Toxicity

**DOI:** 10.1101/2024.10.08.617019

**Authors:** Jack I. Wood, Maciej Dulewicz, Junyue Ge, Katie Stringer, Alicja Szadziewska, Sneha Desai, Srinivas Koutarapu, Haady B. Hajar, Kaj Blennow, Henrik Zetterberg, Damian M. Cummings, Jeffrey N. Savas, Frances A. Edwards, Jörg Hanrieder

## Abstract

It is of critical importance to our understanding of Alzheimer’s disease (AD) pathology to determine how key pathological factors are interconnected and implicated in nerve cell death, clinical symptoms, and disease progression. The formation of extracellular beta-amyloid (Aβ) plaques is the major pathological hallmark of AD and Aβ has been suggested to be a critical inducer of AD, driving disease pathogenesis. Exactly how Aβ plaque formation begins and how ongoing plaque deposition proceeds and initiates subsequent neurotoxic mechanisms is not well understood.

The primary aim of our research is to elucidate the biochemical processes underlying early Aβ plaque formation in brain tissue. We recently introduced a chemical imaging paradigm based on mass spectrometry imaging (MSI) and metabolic isotope labelling to follow stable isotope labelling kinetics (iSILK) in vivo to track the in vivo build-up and deposition of Aβ. Herein, knock-in Aβ mouse models (*App^NL-F^*) that develop Aβ pathology gradually are metabolically labeled with stable isotopes. This chemical imaging approach timestamps amyloid plaques during the period of initial deposition allowing the fate of aggregating Aβ species from before and during the earliest events of plaque pathology through plaque maturation to be tracked. To identify the molecular and cellular response to plaque maturation, we integrated iSILK with single plaque transcriptomics performed on adjacent tissue sections. This enabled changes in gene expression to be tracked as a function of plaque age (as encoded in the Aβ peptide isotopologue pattern) distinct from changes due to the chronological age or pathological severity. This approach identified that plaque age correlates negatively with gene expression patterns associated with synaptic function as early as in 10-month-old animals but persists into 18 months. Finally, we integrated hyperspectral confocal microscopy into our multiomic approach to image amyloid structural isomers, revealing a positive correlation between plaque age and amyloid structural maturity. This analysis identified three categories of plaques, each with a distinct impact on the surrounding microenvironment. Here, we identified that older, more compact plaques were associated with the most significant synapse loss and toxicity.

These data show how isotope-encoded MS imaging can be used to delineate Aβ toxicity dynamics in vivo. Moreover, we show for the first time a functional integration of dynamic MSI, structural plaque imaging and whole genome-wide spatial transcriptomics at the single plaque level. This multiomic approach offers an unprecedented combination of temporal and spatial resolution enabling a description of the earliest events of precipitating amyloid pathology and how Aβ modulates synaptotoxic mechanisms.

## INTRODUCTION

The mechanisms underlying Alzheimer’s disease (AD) pathogenesis are still not fully understood. AD pathology is characterized by extracellular Aβ plaques and intracellular hyperphosphorylated Tau tangles with dementia (Long and Holtzman, 2019). The prevailing model of AD pathogenesis is that changes in Aβ metabolism precipitate a damaging cascade upstream of tau pathology and eventual neurodegeneration (Busche and Hyman, 2020). The relevance of Aβ peptides in AD has seen a recent resurgence as Aβ targeting antibodies lecanemab and donanemab have seen positive outcomes in Phase 3 trials as well as FDA approval (Sims et al., 2023; van Dyck et al., 2023). However, Aβ levels are elevated in AD brains years before synapse loss, circuit dysfunction, neurodegeneration, and impaired cognition (Long and Holtzman, 2019). It is therefore important to capture the disease at the earliest events of Aβ plaque formation. An understanding of Aβ pathogenesis is complicated by the heterogenous presentation of plaque pathology including senile, cored plaques but also diffuse-as well as vascular plaques (Koutarapu et al., 2024; Rohr et al., 2020). At a cellular level, Aβ has been found to alter synaptic vesicle dynamics (Cirrito et al., 2008), where exo- and endocytosis machinery are already altered in the very early stages of Aβ pathology onset (Hark et al., 2021). Clinically, this is in line with the hypothesis that Aβ pathology precipitates several decades before cognitive symptoms occur. Most importantly, these early effects of Aβ aggregation, including synaptic changes, precede the development of mature Aβ plaques that can be detected by PET imaging or by CSF analysis (Cairns et al., 2009; Ikonomovic et al., 2012; Ikonomovic et al., 2008). Exactly how plaques develop over time, the extent of their diversity, and their relation to toxicity or homeostatic response of surrounding neuronal circuits remains unclear (De Strooper and Karran, 2016).

Novel developments within chemical imaging, such as mass spectrometry-based imaging (MSI) together with stable isotope labelling greatly increase our ability to study these events. Using correlative imaging of stable isotope labelling kinetics (iSILK) allows the delineation of plaque formation dynamics in a spatiotemporal manner while maintaining high molecular specificity (Michno et al., 2021).

Herein, we make use of these developments and interface the iSILK approach with both hyperspectral microscopy and spatial transcriptomics. Importantly, the dynamic nature of this approach allows us to track precipitating plaque pathology in the APP^NL-F^ AD mouse model, which mirrors the age-associated and gradual plaque development observed in human Alzheimer’s disease. Together this novel spatial biology approach provides a detailed picture of Aβ-aggregation, plaque formation, and maturation in concert with the associated cellular and molecular response in and around individual plaques at scales not previously possible.

## METHODS

### Animal Experiment

All procedures and experiments on mice were performed at UCL with local ethical approval (06/05/2016) and in agreement with guidelines of the Institutional Animal Care and Use Committee (IACUC) and the Animals (Scientific Procedures) Act 1986. Male and Female APP knock-in mice (*APP^NL-F^*) carrying humanized Aβ sequence, along with the Swedish mutation (KM670/671NL) on exon 16 and the Beyreuther/Iberian mutations (I716F) on exon 17 were used in the study. Transnetyx (Cordova, TN, USA) genotyping services were used to determine the presence of the knock-in genes in breeders.

### Tissue Extraction

Animals were decapitated with the brain rapidly extracted on ice. For the metabolically labelled mice, one hemisphere was snap-frozen directly after isolation using liquid nitrogen (− 150°C)-cooled isopentane. For non-metabolically labelled mice, one hemisphere was snap-frozen directly after isolation on dry ice. In both cases the other hemisphere was drop fixed in 10% formalin (4% PFA) at 4°C overnight, subsequently the formalin was washed out and replaced with 30% sucrose, 0.02% sodium azide in phosphate-buffered saline (PBS) solution for long term storage at 4°C.

### Tissue Sectioning

For correlative GeoMx/ MSI experiments, consecutive, sagittal cryosections were collected at 10μm (slide #A, GeoMx) and 12 μm (slide #B, MSI) were collected from fresh frozen brain tissue on a cryostat microtome (Leica CM 1520, Leica Biosystems, Nussloch, Germany) at - 18°C. Sections were thaw mounted on SuperFrost Plus Slides for the GeoMx DSP platform, with consecutive sections mounted on conductive indium tin oxide (ITO) glass slides (Bruker Daltonics, Bremen, Germany) for MALDI-MSI. All tissue was stored at -80°C.

PFA-fixed brain tissue, collected for time course plaque staining and immunohistochemistry, was sectioned transverse to the long axis of the hippocampus at 30 µm using a frozen sledge microtome (Leica). Free-floating sections were collected and stored in 0.02% sodium azide in PBS at 4°C.

### MALDI-MSI Chemicals

| Reagent | Source |
| --- | --- |
| MouseExpress (15N, 98%) mouse feed kit (15N/14N) | Cambridge Isotope Laboratories (Andover, MA, USA) |
| Acetone (Ac) | Fisher Scientific (Hampton, NH, USA) |
| Acetonitrile (ACN) | Fisher Scientific (Hampton, NH, USA) |
| Absolute ethanol (EtOH) | Fisher Scientific (Hampton, NH, USA) |
| Methanol (MeOH) | Fisher Scientific (Hampton, NH, USA) |
| Glacial Acetic Acid | VWR Chemicals (Radnor, PA, USA) |
| Paraformaldehyde (PFA) | Agar Scientific (Stansted, Essex, United Kingdom) |
| Glutaraldehyde | Agar Scientific (Stansted, Essex, United Kingdom) |
| Sodium cacodylate buffer | Agar Scientific (Stansted, Essex, United Kingdom) |
| Osmium tetroxide | Agar Scientific (Stansted, Essex, United Kingdom) |
| Agar 100 resin | Agar Scientific (Stansted, Essex, United Kingdom) |
| Formic acid (FA) | Honeywell Fluka, Germany |
| Trifluoroacetic acid (TFA) | Honeywell Fluka, Germany |
| 2,5-Dihydroxyacetophenone (2,5-DHAP) | Sigma-Aldrich (St. Louis, MO, USA) |
| TissueTek optimal cutting temperature (OCT) compound | Sakura Finetek (AJ Alphen aan den Rijn, Netherlands) |
| Indium tin oxide (ITO)-coated conductive glass slides | Bruker Daltonics (Bremen, Germany) |
| Peptide calibration standard | Bruker Daltonics (Bremen, Germany) |
| Protein calibration standard | Bruker Daltonics (Bremen, Germany) |
| PELCO copper grids | Ted Pella (Redding, CA, USA). |

### Matrix deposition for MALDI-MSI

Fresh frozen tissue sections on ITO-coated glass slides were fixed in 100% ethanol for 60 seconds, followed by 70% ethanol for 30 seconds. Lipids were removed by immersing the sections in Carnoy’s solution (6:3:1 ethanol/chloroform/acetic acid) for 110 seconds, followed by subsequent washes in 100% ethanol for 15 seconds, 0.2% trifluoroacetic acid in water for 60 seconds, and 100% ethanol for 15 seconds. The tissues then underwent peptide retrieval using formic acid vapor for 20 minutes. To facilitate peptide ionization and desorption, a matrix solution (15 mg/ml 2,5-dihydroxyacetophenone in 70% acetonitrile, 2% acetic acid, 2% trifluoroacetic acid) was applied using a TM sprayer (HTX Technologies, NC, USA) and recrystallized in 5% methanol vapor at 85°C for 3 minutes.

### MALDI-MSI Data Acquisition

MALDI-MSI was performed using a MALDI-time-of-flight (TOF) instrument (RapifleX, Bruker Daltonics) equipped with a scanning Smartbeam 3D laser. The acquisition was conducted at a pixel size of 10 μm with 200 shots per pixel at 90% power with a shot frequency of 10 kHz Linear positive mode was used due to higher sensitivity at high spatial resolution. Spectra were collected within the mass range of 1500–6000 m/z (mass resolution: m/Δm=1000 (FWHM) at m/z 4515).

For accurate detection of Aβ, the system was externally calibrated by spotting a mixture of peptide standard II and protein standard I (Bruker Daltonics). Total ion current normalization for each ROI was performed using FlexImaging 5.1 software (Bruker Daltonics). Binning analysis, including the calculation of peak width and amplitudes per average spectra, was conducted using the peak analyzer function in Origin (v.8.1; OriginLab, USA).

### Nanostring GeoMx Digital Spatial Profiler

Slides were processed by the Nanostring Technology Access Programme (TAP; Seattle, WA, USA) following the standardized GeoMx protocol for fresh frozen samples.

#### Slide Preparation

The 10 µm cryosections were fixed in 10% formalin followed by ascending concentrations of EtOH. Slides were then immersed in 1X Tris EDTA at 100°C for 15 minutes in a pressure cooker. After washing in PBS, RNA targets were exposed to 1ug/µl of Proteinase K in PBS at 37°C for 15 minutes. Tissues were post-fixed in 10% formalin for 5 minutes at RT followed by 2* 5-minute washes in NBF Stop Buffer at RT (see GeoMx manual). Tissues were subsequently incubated in whole transcriptome RNA probes overnight in a hybridization chamber held at 37°C. Off-target probes were washed out in 2* Stringent Solution (see GeoMx manual) for 25 minutes at RT. Nonspecific binding was blocked using Buffer W (Nanostring propriety blocking buffer) followed by incubation in conjugated antibodies in Buffer W for 2 hours at RT (mouse anti-GFAP Alexa-Fluor 488 conjugate (Invitrogen, #53-9892-82), mouse anti-Aβ40/42 Alexa Fluor 594 conjugate (Nanostring, #121301306). After which, nuclei were counterstained for 1 hour at RT (SYTO 13, #S7575). After a final wash in 2x SSC buffer, slides were placed into the GeoMx machine.

#### ROI Selection

MALDI MSI images of consecutive sections were overlayed on the fluorescent image output from the GeoMx. Plaques spanning both sections were identified, and a circular region of interest (ROI) three times the radius of the plaque was centered around the plaque’s midpoint. All probes regardless of cell type were extracted per ROI.

##### Library preparation and readout

DNA barcode oligomers were aspirated into well plates whereby the aspirate from one plaque corresponds to one well. The well plates were then dried and rehydrated in 10µl nuclease-free water and PCR cycled with GeoMx SeqCode master mix and SeqCode primers (Nanostring Technologies). Finally, the PCR product was purified with AMPure beads. Plates were subsequently sequenced on an Illumina NextSeq2000. FASTQ files were converted to .dcc with the GeoMx NGS Pipeline GUI.

#### GeoMx Data Analysis

The Quality control (QC) analysis for NGS and Biological Probe QC was conducted utilizing the default GeoMX DSP parameters (GeoMX NGS Pipeline 2.3.4). All further analyses were carried out in R [R Core Team, Vienna Austria, https://www.R-project.org/]. Probes were excluded from analysis if counts were too low or failed the Grubbs outlier test according to NanoString guidelines. Nine ROIs in the 10-month-old group and six ROIs in the 18-month-old group were excluded from further analysis due to ambiguous spot morphology >1 plaque or very low intensity in MALDI results. After data filtering, the geometric mean was calculated for targets in each AOI. QC results were compared with those from the standard NanoString pipeline, showing generally consistent outcomes in outlier detection. Any minor discrepancies are likely attributable to slight differences in the test parameters. Quantile normalization was applied using the normalize.quantiles function provided by the preprocessCore R package. Pearson’s correlation was run between quantile normalised gene expression and peak data from MALDI MSI. The local intersection with the fitted baseline was assessed for each Aβ 1-42 peak derived from MALDI imaging data (Figure 3). The correlation results were divided into positive and negative, considering age (10-month-old positively/negatively and 18-month-old positively/negatively correlated). Gene over-representation enrichment (ORA) analysis was performed for each group separately, including the three different Gene Ontology (GO) terms related to Biological Processes (BP), Cellular Component (CC) and Molecular Functions (MF). For the ORA analysis, we applied a p-value and q-value cutoff of 0.05, using the entire gene set as a background. This analysis was conducted using the two R packages, geneset and genekitr. Data visualizations were carried out using in-house functions and modification of already existing functions, as a part of packages: GOplot, ClusterProfiler, tidyr.

### Immunohistochemistry and LCO staining

Free-floating 30 µm sections underwent antigen retrieval in 10mM PH9 Tris-EDTA for 30 minutes at 80°C. Subsequently, sections were permeabilized in 0.3%Triton X-100 in PBS (PBST) for 3* 10 minutes at RT before non-specific binding was blocked in 3% goat serum in PBST (Blocking solution) for 1 hour at RT. Sections were incubated with primary antibodies diluted in blocking serum at 4°C overnight (1:500 mouse anti-Aβ (6E10, #SIG-39320), 1:500 rabbit anti-Aβ (#700254), 1:200 chicken anti-HOMER1 (#160001), 1:500 rat anti-LAMP1 (#ab25245)). Sections were washed in PBST 3* 10 minutes at RT followed by incubation in secondary antibodies diluted in blocking solution for 2 hours at RT (1:500 goat anti-mouse AF594 (#A11032), 1:500 goat anti-rabbit AF594 (#A11037), 1:500 goat anti-chicken AF647 (#A-21449), 1:500 Donkey anti-rat AF594 (#A-21209)). Nuclei were counterstained with 4ʹ,6-diamidino-2-phenylindole (#ab228549) for 5 minutes at RT.

For structural amyloid analysis, tissues were stained with two luminescent conjugated oligothiophene (LCO) dyes, tetra-(q) and heptameric (h-) formyl thiophene acid (FTAA). For linear unmixing analysis after antibody staining, sections were subsequently incubated for 30 minutes at RT with 1:500 q-FTAA and 1:1500 h-FTAA.

For LCO MALDI correlation analysis, cryo-sections were fixed sequentially in a gradient concentration of ice-cold 95%, 70% ethanol, and 1× phosphate-buffered saline (PBS) at room temperature followed by incubations in q-FTAA, 2.4 μM in Milli-Q water and h-FTAA, 0.77 μM in Milli-Q water) in the dark for 25 minutes.

### Fluorescent Microscopy

All imaging was performed on the Zeiss LSM780 confocal microscope. The imaging of double LCO-stained tissues was performed in hyperspectral mode using the 32-channel GaAsP spectral detector. The dyes were stimulated using a 35nW, 458nm argon laser. For LCO correlation analysis, continuous emission spectra were obtained in lambda mode. For plaque-type classification, linear unmixing mode utilized reference spectra of q-FTAA and h-FTAA to isolate the corresponding signals as individual channels.

For LCO correlation analysis individual amyloid plaques were randomly selected and images as z-stacks (20 stacks per plaque, 1-μm apart) under a 20X air objective. The 500:580 emission ratios were collected using Zen Black 2.3 software.

In the linear unmixing analysis, LAMP1, Aβ, HOMER1, q-FTAA and h-FTAA images were taken as z-stacked (3 µm apart) tile scans of whole hippocampus using a 20X air objective. LAMP1, Aβ and HOMER1 were imaged in channel mode using constant light, gain and exposure settings matched to the fluorophore.

## Image Analysis

### LCO and MALDI MSI correlation analysis

Z-stacked images of individual plaques were analyzed in lambda mode, selecting the plane that showed the most pronounced 500nm shift for further examination. Within this plane, a small circle, just a few pixels in diameter, was placed on the area of the plaque exhibiting the highest 500nm shift, indicating the core. At this location, intensities at both 500nm and 580nm were recorded for analysis.

For the MALDI imaging data acquired in reflector mode, regions of interest (ROIs) were annotated, and total ion current normalized average spectra of the annotated ROIs were exported as .csv files in flexImaging. This was followed by a binning analysis for data reduction. Here, all ROI data were imported into Origin (v 8.1 OriginLab, Northampton, MA, USA) and the Peak Analyzer function in Origin was used to determine the isotopic peak area of Aβ 1-42 within each ROI spectra. The ratio of the 4th to the 3rd isotopic peak areas of Aβ 1-42 was used to determine the relative age of the plaque.

A correlation analysis was then conducted between the 500/580 nm ratio in hyperspectral data and the corresponding 4th/3rd isotopic peak area ratio in MALDI imaging data. 3-5 plaques from 3 animals were selected for this correlation analysis.

### Protein intensity towards plaque type

Image analysis was carried out using a set of custom macros for ImageJ. In short, the Z-stack tile scans were projected to a single plane based on standard deviation. Regions of interest (ROIs) were then drawn around each hippocampus to restrict further analysis to this area. Plaques were subsequently thresholded and categorized based on the positivity of both Aβ and LCO dyes. Radiating rings from each plaque were created at 10 µm increments extending up to 30 µm with any overlapping areas removed from the analysis. Mean grey values for each protein of interest at each increment were averaged per plaque type for each hippocampal image. 2-3 hippocampal images were averaged for each animal.

Due to technical difficulties, LAMP1 analysis was conducted in two stages: The first stage involved assessing LAMP1 in plaques that were both Aβ and LCO positive (specifically h-FTAA only) and in plaques that were Aβ positive but LCO-negative. The second stage focused on plaques positive for both h-FTAA and q-FTAA, as well as plaques positive for h-FTAA but negative for q-FTAA. In contrast, HOMER1 analysis with Aβ, h-FTAA, and q-FTAA staining, was completed in a single batch.

### Statistical Analysis

Statistical analysis was performed in either GraphPad Prism 9 or RStudio. Post hoc analysis using Sidak’s correction was applied only when a statistically significant interaction was detected. For the analysis of center versus periphery (Figure 1), a linear mixed-effects model was employed to account for paired measurements within plaques and to account for variability between biological samples. In all statistical tests biological replicates were considered appropriately to avoid pseudoreplication.

**Figure 1.**
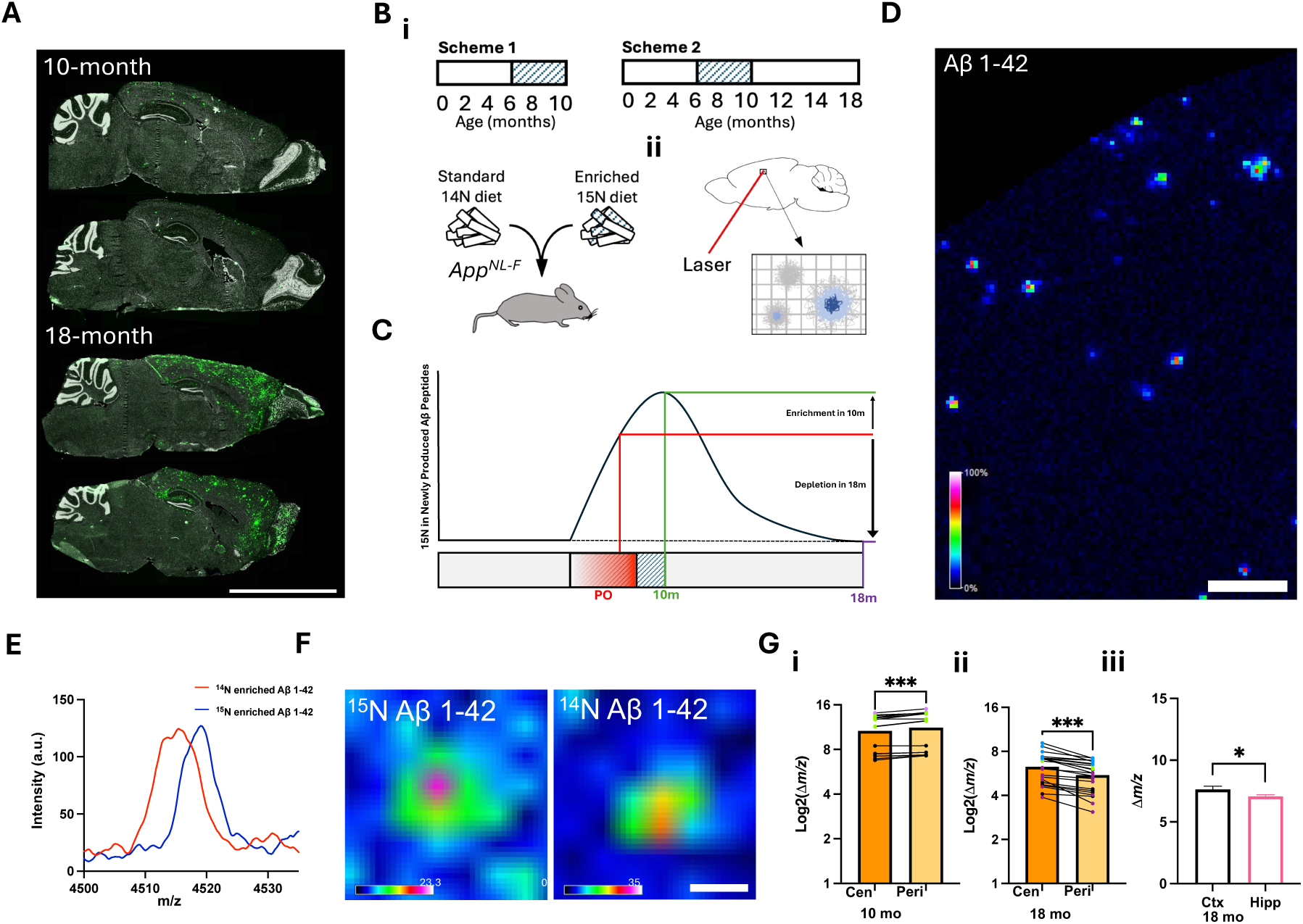
iSILK delineates spatial and structural patterns of plaque formation and maturation. (A) Representative sagittal images of plaque load (green) of 10- and 18-month-old APP^NL-F^ mice (B) Pulse-chase iSILK paradigm (i) feeding schemes for ^15^N enriched diet, pulse period between 6-10 months of age. PO indicates plaque onset, with initial hippocampal plaques appearing between 6-9 months (iii) MALDI MSI scheme detailing the collection of Aβ peptides from defined rasters. (C) Predicted ^15^N enrichment of newly synthesized Aβ peptides over the course of the pulse-chase feeding paradigm. (D) MALDI MSI produced single ion image of Aβ1-42 in a section of temporal cortex. Scale bar 200μm (E) Example spectra from MALDI MSI showing the Aβ1-42 m/z peak. (F) Representative MALDI MSI image of ^15^N and ^14^N enriched plaques. Scale bar 20μm (G) Aβ1-42 mass analysis comparing (i) the plaque centre (Cen) vs. the periphery (Peri) in 10-month-old mice, (ii) in 18-month-old mice, and (iii) differences between cortex and hippocampus. (i, ii) Linear Mixed Model, animal indicated by point color, (iii) Paired t-test. Data presented as mean ± SEM. Significance levels: *** P<0.001, **P<0.01; *P<0.05.

## Results

### 1. iSILK delineates spatial and structural patterns of plaque formation and maturation

Initial amyloid aggregation and subsequent plaque formation are dynamic processes, and difficult to capture whilst tracking across chronological ageing. We previously demonstrated the potential of stable isotope labelling and MSI to follow Aβ plaque formation in the same-aged *App^NL-G-F^* mice (Michno et al., 2021). While these initial experiments were key for establishing the technique and its feasibility, there are limitations with respect to the *App^NL-G-F^* model whereby the Arctic mutation of APP alters the chemical properties of Aβ leading to an increased aggregation propensity not seen in sporadic AD. This results in a rapid plaque onset with a pronounced formation of cored deposits, lacking the range of plaque isoforms seen in the human disease (Saito et al., 2014). Therefore, by building on the previous findings from *App^NL-G-F^* mice, we aimed to follow Aβ aggregation in the more slowly progressing plaque pathology of *App^NL-F^* mice. This model provides a more physiological presentation of amyloid aggregation due to a gradual increase in Aβ. Plaques appear infrequently as early as 6 months with very low plaque levels persisting until 9 months, increasing thereafter to reach a plateau by 18 months (Figure 1A, Benitez et al., 2021; Saito et al., 2014). This gradual increase in plaque pathology with age more closely resembles the human disease.

For this study, *App^NL-F^* mice were labelled with ^15^N spirulina diet from age 6-10 months (PULSE period, 4 months, Fig. 1Bi) and either culled immediately at 10 months old (Figure 1Bi, Scheme 1), or after an 8-month CHASE (washout) at 18 months old (Figure 1Bi, Scheme 2). Here, *App^NL-F^* mice at 10 and 18 months of age represent early and established stages of plaque pathology, respectively (Figure 1A).

For scheme 1, continuous labeling was used to identify whether the labelling scheme was implemented in time to ensure that secreted Aβ peptides were labelled prior to extracellular deposition and precipitation in plaques (Saito et al., 2014). MALDI MSI was then used to determine stable isotope enrichment in plaque-associated, deposited Aβ peptides in situ (Figure 1Bii and 1C). This approach enables us to image plaques by spatially measuring Aβ1-42 species (Figure 1D) and correlating the isotope enrichment of the deposited Aβ peptides with the different plaque structures such as the center and periphery, thereby investigating the temporal development of these features during plaque formation.

At 10 months of age, the sparse deposition of Aβ plaques in the cortex contained solely Aβ1-42. Importantly, it was observed that all Aβ1-42 peptides detected showed ^15^N incorporation (Figure S1). This ensured that the onset of the labelling scheme at 6 months old was sufficient to incorporate the ^15^N label into APP prior to plaque pathology onset, which is typically observed at 9 months in this model. After establishing the feasibility of the iSILK approach in *App^NL-F^* mice, we then set out to interrogate deposition dynamics within and across plaques and brain regions.

First, to address the question of intra-plaque heterogeneity (center vs. periphery), the relative ^15^N content of these plaque structures was evaluated. For this, we isolated plaque ROI spectral data of hippocampal and cortical deposits (Figure 1). We compared the isotopologue signature of plaque-associated Aβ1-42 that encodes the degree of labelling, as indicated by the shift in mass (Figure 1). For an unbiased, quantitative comparison of label incorporation, we performed curve analytics of the isotope envelope for a plaque-associated signal that in turn encodes the degree of label incorporation. Here, to estimate the shift in m/z value as caused by the ^15^N label and hence the difference in label content, we compared the centroid of the curve fitted to the Aβ1-42 isotopologue pattern, attributed to the comparably low mass resolution in linear mode (Figure 1E). These tradeoffs with linear mode were needed to ensure sensitive peptide detection at high spatial resolution.

Based on the labelling scheme, this shift in the isotopologue signature indicates earlier or later deposition. Here, plaques in 10-month-old *App^NL-F^* mice (PULSE age 6-10 months, no CHASE) displayed lower ^15^N content for Aβ1-42 in the center compared with the periphery (Figure 1C, 1F and 1Gi, P<0.01). This suggests that previously deposited Aβ1-42 species have less ^15^N at the center, while later secreted Aβ1-42 deposits at the periphery. In 18-month-old animals (Scheme 2, PULSE age 6-10 months, 8-month CHASE), the pattern was reversed due to the significant deposition of unlabeled Aβ1-42 during the CHASE period (Figure 1C and 1Gii, P<0.05).

Both experiments suggest indeed that Aβ pathology in *App^NL-F^* mice appears to be initiated through the formation of small compact deposits constituting the center of the forming plaque followed by homogenous plaque-wide deposition during the plaque growth phase.

While these first analyses identified deposition patterns within plaques, iSILK further allows us to delineate plaque formation sequences across different brain regions. This is important as it allows for a description of how plaque pathology spreads throughout the brain and in what sequence it affects vulnerable brain regions. Similar to the plaque heterogeneity analyses, isotope incorporation was determined from the isotopologue patterns of Aβ 1-42 in plaque features detected in the cortex and hippocampus as described above. The results show that the degree of isotope content, as expressed as the centroid of the fitted isotopologue distribution curve, was higher in cortical plaques compared to hippocampal plaques (Figure 1Giii). This indicates that the cortex is the primary location of initial plaque formation, which is well in line with immunohistochemical studies described for this model as well as previous iSILK data obtained for *App^NL-G-F^*mice (Michno et al., 2021).

### 2. iSILK-guided spatial transcriptomics shows changes in synaptic-, metabolic-, and immune-associated gene expression with plaque age

Utilizing the iSILK technique to track plaque age, we conducted single plaque spatial transcriptomics to monitor gene expression changes associated with plaque age, independent of the mouse’s chronological age. Spatial transcriptomics was selected over other sequencing techniques due to its ability to target plaque-specific gene expression changes, offering a more spatially resolved technique for AD pathology-associated alterations compared to the more commonly used RNA sequencing methods (Smith et al., 2024, Figure S2). The regions of interest (ROIs) chosen on the spatial transcriptomics platform were selected by identifying double positive (MALDI and IHC) plaques across two serial sections, with the MALDI MSI images imported and overlaid onto the IHC image for precise mapping (Figure 2A, 2B, and Table S1). As plaques were selected across serial sections, the plaques analyzed were all large and therefore plaque size showed no significant correlation with plaque age (P = 0.25, r=0.29).

**Figure 2.**
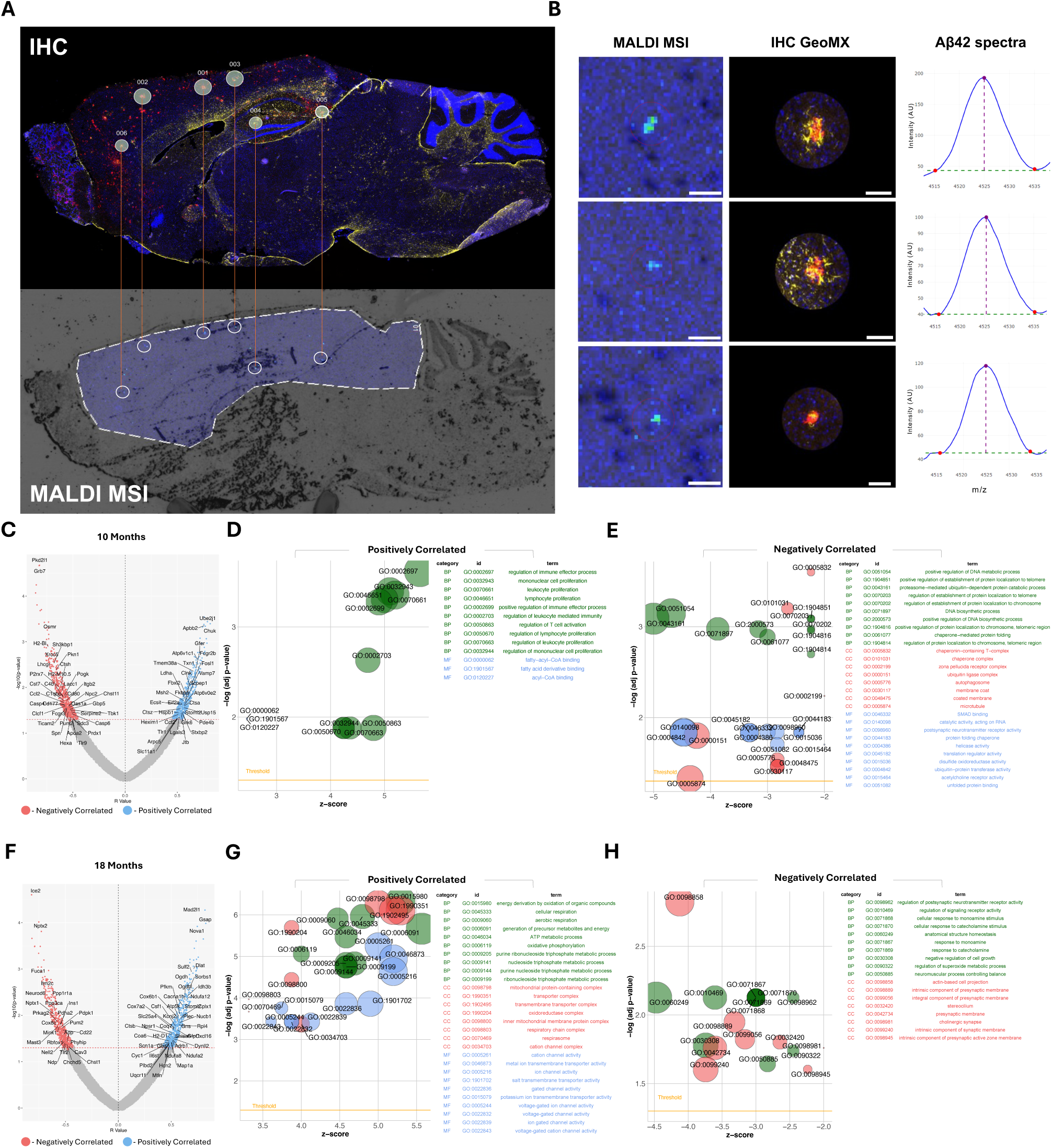
iSILK guided spatial transcriptomics shows changes in synaptic-, metabolic-, and immune-associated gene expression with plaque age. (A) Matching of Aβ plaques on consecutive sections for immunohistochemistry (IHC) and MALDI MSI imaging. (B) Aβ signals of single plaques across MALDI MSI and IHC images, including m/z spectra with peak and centroid calculations. Scale bar: 100μm. (C) Volcano plot for gene expression correlations with increasing plaque age in 10-month-old *App^NL-F^* mice (Scheme 1). (D, E) Gene ontology analyses of positively (D) and negatively (E) correlated genes in 10-month-old mice, presented in bubble plots. (F) Volcano plot for gene expression correlations with increasing plaque age in 18-month-old APP^NL-F^ mice. (Scheme 2). (G, H) Gene ontology analyses of positively (G) and negatively (H) correlated genes in 10-month-old mice, presented in bubble plots.

To identify associations between the plaque age (MALDI iSILK) and whole transcriptome-wide gene expression, we performed correlation analysis between peak centroid and quantile normalized counts for all genes. Volcano plots for both 10-month (Figure 2C) and 18-month-old (Figure 2F) mice demonstrate significant up- and down-regulation of genes associated with plaque age. Correlation statistics for all genes whole transcriptome can be searched at: https://hanriederlab.shinyapps.io/PlaqueAgeTranscriptomics/.

Functional annotation of correlated genes at 10 months of age (Scheme 1) shows that older plaques are associated with increased expression of genes related to immune regulation and immune cell proliferation (*H2-T24*, *H2-M10.5*, *H2-Bl*, *Itgb2*, *Apoa2*, Figure 2D and S3). In contrast, genes related to synaptic activity, particularly acetylcholine-associated (*Chrnb1*, *Chrnd*, *Chrm3*, *Chrm2*, *Gria4*) and protein trafficking were negatively associated with plaque age (*Ubxn7*, *Igfbp3*, Stoml2, Figure 2E and S3). In 18-month-old animals (Scheme 2), the analysis reveals that older plaques are linked to an increase in genes involved in metabolic processes (*Atp5a1*, *Atp5k*, *Cox7a2*, *Coa6*, *Coq7*) and channel activity (*Kcnj2*, *Kcna2, Scn1a*, *Scn2b*, *Gria3*, Figure 2G and S3). In contrast, older plaques correlated with a concomitant decrease in genes predominantly associated with synapses and neurotransmitter receptors (*Shisa6*, *Chrna4*, *Drd1*, *Nptx2,* Figure S3).

Using synaptic hub genes (Williams et al., 2021) and disease-associated genes (Habib et al., 2020), we compared expression profiles between previously generated spatial transcriptomics data and RNA-seq results from bulk hippocampal tissue analysis (Wood et al., 2022, Figure S2). While spatial transcriptomics effectively captured expression changes associated with Alzheimer’s pathology, bulk RNAseq was less sensitive in detecting these differences. Additionally, incorporating plaque age, as enabled by iSILK, reveals further age-related changes that are overlooked when only considering plaques at a given chronological age (Wood et al., 2022, Figure S2). This underscores the significant advantages of spatial transcriptomics in addressing diseases with strong spatial characteristics, such as AD.

In summary, at 10 months, older plaques are associated with increased immune-related gene expression, while at 18 months, they show higher transcription of metabolism-related genes. In both age groups, plaque aging is consistently linked to a decline in synapse-related gene expression.

### 3. Amyloid plaque maturation is characterized by continuous fibrilization with age

In addition to changes in gene expression, we also aimed to investigate the structural maturation of plaques as they age. Therefore, we complemented the iSILK MSI experiments with fluorescent microscopy using the structure-specific luminescent conjugated oligothiophene (LCO) amyloid probes (Almstedt et al., 2009; Nilsson et al., 2007, Figure 3A). LCO staining paired with hyperspectral confocal microscopy was performed on consecutive sections allowing for the identification of morphologically heterogenous amyloid structures within the Aβ plaques. This is enabled by the difference in affinity of the two LCO probes, q-FTAA and h-FTAA, towards amyloid aggregates. Specifically, q-FTAA preferentially binds to mature and compact beta-pleated aggregates, while h-FTAA binds to less compact, yet still beta-pleated aggregates. Due to their different emission profiles, the LCO probes can be spatially delineated using hyperspectral fluorescent microscopy. Here, the ratio of the LCO maxima (500nm for q-FTAA / 580nm for h-FTAA) is used to express preferential binding of either of the two LCO probes used, whereby an increase in 500nm intensity is indicative of increased q-FTAA binding and therefore, increased structural maturity of the amyloid fibrils (Figure 3B). First, we focused on 18-month-old *App^NL-F^* animals (Scheme 2, Figure 1Bi) and compared plaques in the cortex to those in the hippocampus. Similar to the iSILK data showing that cortical plaques are older than those in the hippocampus, our results demonstrate that cortical plaques also exhibit higher 500/580 nm ratios at their centers, suggesting that they are more aggregated (Figure 3C). To further test this hypothesis, we performed a single plaque correlation analysis of correlative iSILK/LCO data collected from plaques detected across two sequential tissue sections. The results indeed, showed a positive correlation (R 0.63-0.89, p<0.05, Figure 3E) of label incorporation and LCO emission ratio (Figure 3D). Together these data are the first of its kind showing a direct correlation of differentially aged, individual plaques with biophysical measures of amyloid aggregation within individual animals. These results confirm that in these mice, plaques appear to continuously mature in terms of amyloid fibrillization at the plaque core.

**Figure 3.**
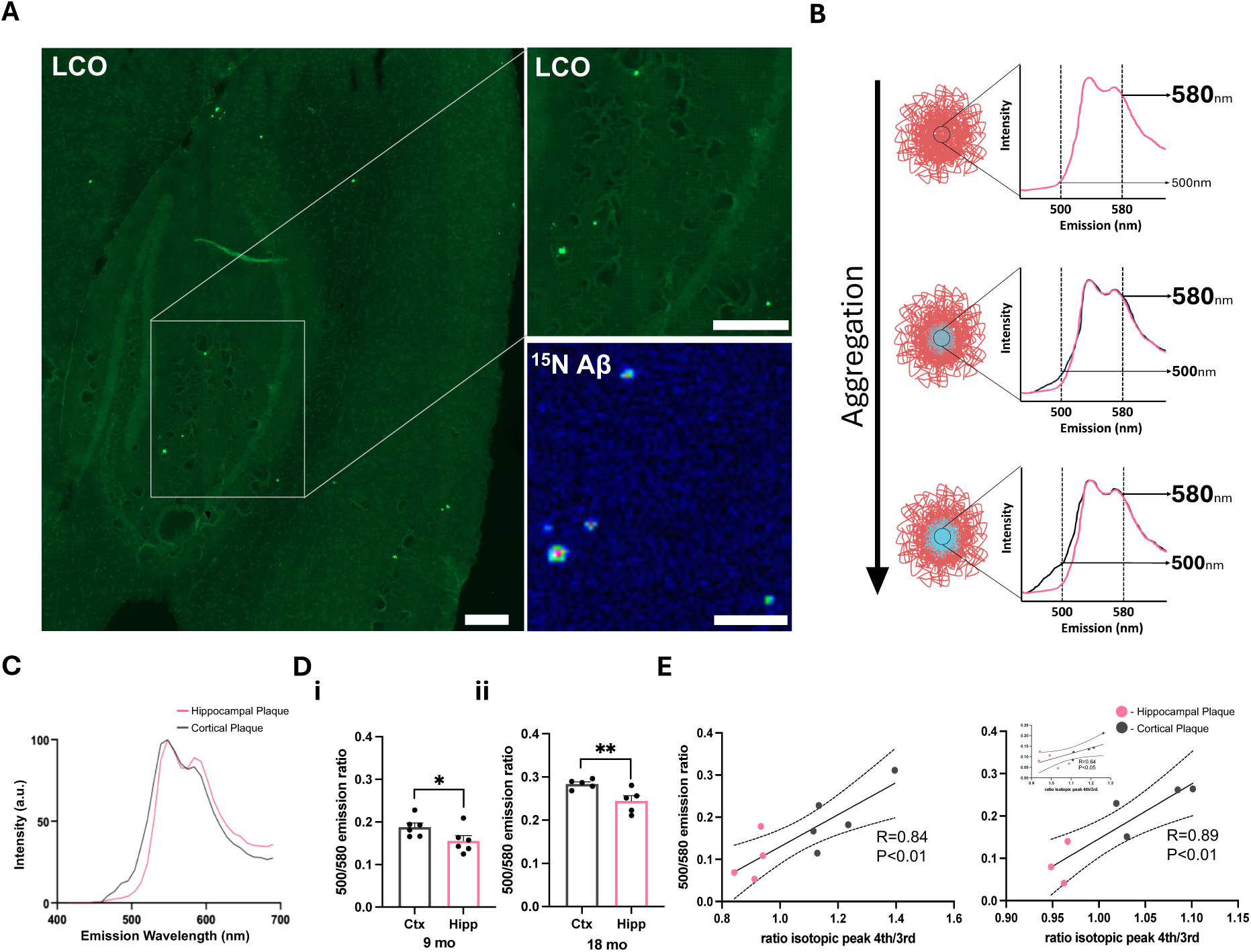
Amyloid plaque maturation is characterized by continuous fibrilization with age. (A) Representative images demonstrating plaque detection using LCO staining and MALDI MSI. Scale bar 200μm (B) Schematic illustrating increasing 500nm intensity resulting from q-HFTAA binding as plaques aggregate. (C) Averaged mass spectra comparing hippocampal and cortical plaques. (D) Analysis of the emission ratio, with the 500nm q-HFTAA peak against the 580nm h-HFTAA peak, in cortex vs. hippocampus for 9-month-old (i) and 18-month-old (ii) mice. Paired t-test (E) Pearsons’s correlation analysis of the 500/580nm ratio with the Aβ1-42 isotopic peak ratio for three 18-month-old mice. Data presented as mean ± SEM. Significance levels: **P<0.01; *P<0.05.

### 4. Differences in Synapse Loss and Toxicity Revealed by LCO-Defined Plaque Types

As amyloid aggregation / LCO probe binding is a factor of plaque age, we adapted our hyperspectral imaging approach to categorize amyloid plaques depending on the presence of q-FTAA and h-FTAA. Linear unmixing, implemented in hyperspectral mode, utilized reference spectra of pure q-FTAA and h-FTAA to separate the probes into distinct channels (Figure 4A). Additionally, an Aβ42 antibody served as a general Aβ marker. We focused exclusively on 18-month-old animals, as the 10-month-old group presented too few hippocampal plaques to yield a sufficient dataset. We identified three populations of structurally distinct plaque types depending on the positivity of the LCO probes: Aβ+h+q+, Aβ+h+q-, and Aβ+h-q-, here arranged in descending order of aggregation maturity (Figure 4B). In the paired LCO/iSILK results above, we established that continuous plaque aging is associated with increased q+ staining i.e. amyloid fibrilization (Figure 3E). This suggests that Aβ+q+h+, Aβ+q+h-, and Aβ+q-h-represent distinct stages of plaque maturation. Therefore, we were able to study plaque ages dissected from chronological age within the same animal using fluorescent microscopy-based correlated LCO imaging and immunohistochemistry.

**Figure 4.**
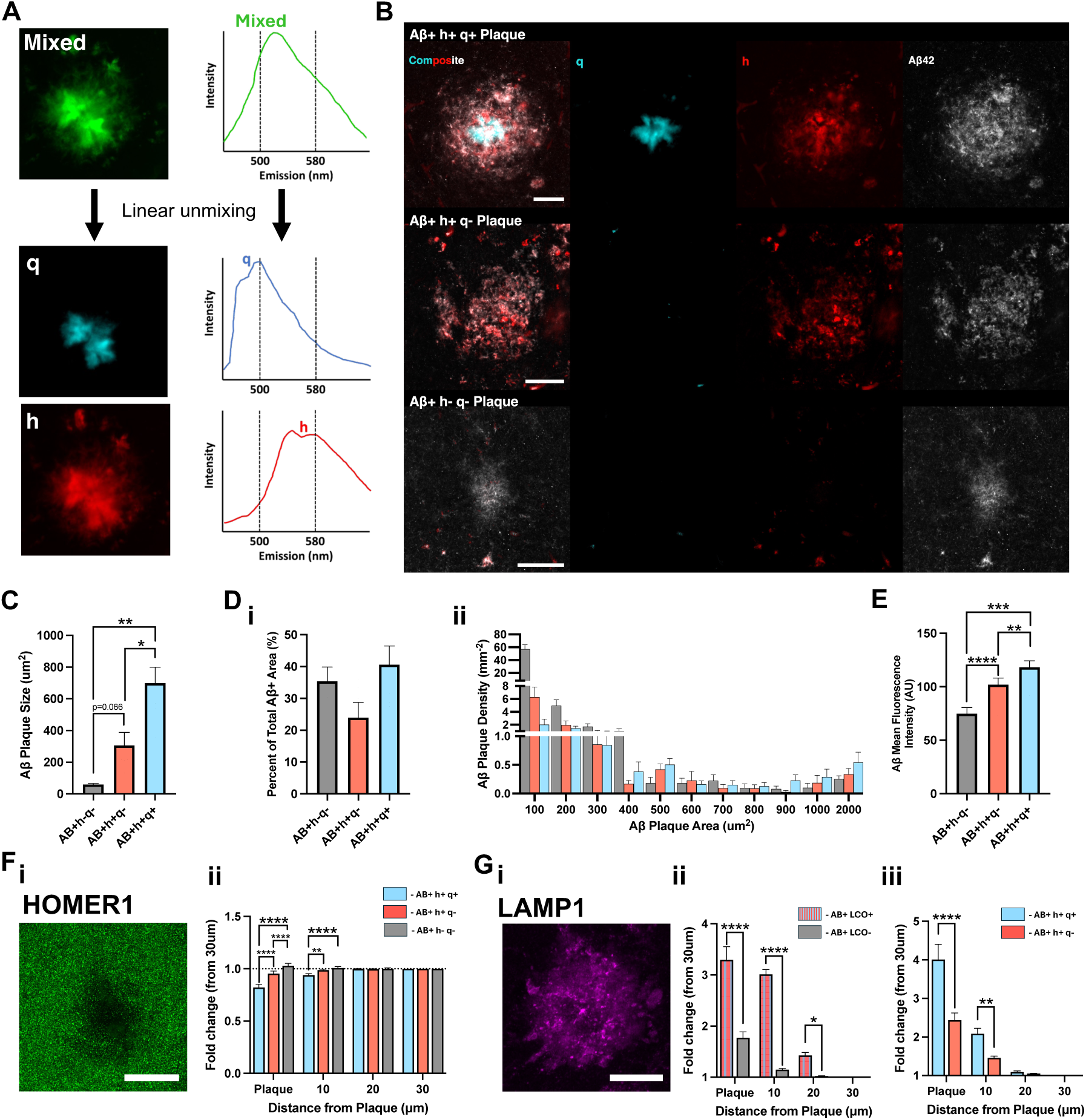
Differences in Synapse Loss and Toxicity Revealed by LCO-Defined Plaque Types. (A) Schematic illustrating the linear unmixing of mixed LCO signal into isolated q-HFTAA and h-HFTAA channels. (B) Representative images of three LCO defined plaque types. (C) Average area covered by individual plaques. (D) (i) Percentage of total Aβ area covered by each plaque types. (ii) Histogram distribution of plaque areas. (E) (Average Aβ fluorescence intensity of plaques. (F) (i) Image illustrating HOMER1 loss associated with plaques. (ii) Analysis of protein fluorescence intensity relative to LCO-defined plaque types. (G) (i) Image showing increased LAMP1 expression at plaques. (ii, iii) Quantification of protein fluorescence intensity relative to LCO-defined plaque types. Statistical Analysis: (C, D, E) One-way paired ANOVA, n=6. (F, G) Two-way repeated measures ANOVA. (F: n=6, G(i): n=7, G(ii): n=4). Data in panels (C, D, E) are plotted as paired data, while panels (F, G) data are presented as mean ± SEM. Post-hoc Tukey’s test corrections applied. Significance levels: ****P<0.0001; ***P<0.001; **P<0.01; *P<0.05. Scale bar 20μm.

On average, Aβ+h+q+ plaques were the largest in area accounting for 41% of Aβ-positive regions in the hippocampus, followed by Aβ+h+q-plaques, which occupied 24%. A notable finding was the detection of many small, LCO-negative (Aβ+h-q-) plaques in the hippocampus, which constituted 35% of the plaque-positive area (Figure 4C & 4D). Assuming increased Aβ compaction means increased Aβ peptides per pixel and therefore higher fluorescence intensity, we measured the average intensity per plaque type. As hypothesized, Aβ fluorescence intensity increased with increased plaque aggregation (Figure 4E). Small immature plaques of low Aβ intensity are, however, difficult to capture in MALDI MSI, particularly spanning across two sections for use in correlative GeoMX.

When examining these plaque types over the lifespan of the mice, we observed that Aβ+h+q- and Aβ+h+q+ plaques together make up 90% of the Aβ-positive area in the hippocampus of younger, 9-month-old *App^NL-F^*mice. This suggests that small fibrillized plaques are likely the initial precipitating plaque type (Figure S4). By 14 months, the proportion of Aβ-positive area occupied by Aβ+h+q-plaques rose to 26% (53% of plaques by number), increasing further to 36% (78% of plaques by number) by 18 months (Figure S4). This indicates a significant accumulation of non-pleated plaques as the mice age.

Together these findings support the notion that at pathology onset, plaques form at small cores which continuously mature through progressing amyloid deposition. We consequently focus on Aβ+h+q+ that are continuously detected through all stages of progressing plaque pathology. Considering the observation above that increasing plaque age is associated with changes in transcription of synaptic genes (Figure 3), we measured synapse loss and toxicity surrounding the plaque types using antibodies against HOMER1 and LAMP1 respectively. Synapse loss correlated with increased plaque aggregation, with Aβ+h+q+ plaques demonstrating the greatest reduction in HOMER1 signal (Figure 4F). This corresponded to higher levels of dystrophic neurites (LAMP1) around more aggregated plaque types (Figure 4G).

## Discussion

Most studies attempting to understand the development of AD and neurodegeneration primarily consider factors such as chronological age and pathological severity. However, a detailed understanding of AD development still lacks a single plaque evolution-related perspective. For instance, it is unclear whether newly formed plaques in the same individual exert a different influence on surrounding tissues compared to older plaques. Furthermore, it remains an important question in AD research why Aβ plaques have a poor correlation with cognitive decline despite the amyloid hypothesis suggesting it as the disease trigger (Morris et al., 2014). This gap in understanding is further emphasized by a population of people who live dementia-free lives but upon post-mortem have a brain populated with plaques (CU-AP individuals, Murray and Dickson, 2014; Serrano-Pozo et al., 2011). Interestingly, the brains of these patients are characterized by an abundance of diffuse plaques, suggested to be an immature state that exerts little toxicity to its surroundings. These findings suggest a close relationship between plaque age, structural maturity, and neurotoxicity. In this study, using a novel multiomic iSILK-driven approach, we demonstrate exactly this. In older mice, older plaques are associated with greater neurotoxicity, more extensive loss of synapses, and higher structural maturity compared to newly formed plaques.

Traditional bulk sequencing methods required mouse models of Alzheimer’s disease to develop exaggerated levels of plaque pathology in order to minimize non-pathological tissue to capture plaque-induced transcriptomic changes. Therefore, more disease-relevant Alzheimer’s disease models such as *App^NL-F^* mice, that develop plaques gradually, with onset in older ages, and without APP overproduction were unable to produce sufficient plaque loads to influence bulk sequencing results (Benitez et al., 2021; Ghosal et al., 2009; Sasaguri et al., 2017, Figure S2). While our previous work, along with other studies, has leveraged recent advancements in spatial transcriptomics to demonstrate plaque-specific alterations in gene expression (Chen et al., 2020; Choi et al., 2023; Mallach et al., 2024; Matarin et al., 2015; Wood et al., 2022) these studies often combine plaque regions and therefore do not consider the heterogeneity of plaque types and maturity. This raises questions about the differential effects of plaque heterogeneity on surrounding tissue. For the first time, we have integrated spatial transcriptomics with an iSILK MALDI MSI paradigm to provide insights into how gene expression evolves in conjunction with plaque maturation at the single plaque level. We observed a negative correlation between synaptic-associated genes and plaque age at both 10 months (Scheme 1) and 18 months (Scheme 2), indicating that while the loss of synaptic function or synaptic puncta is associated with increasing plaque age, it still occurs at both early and late chronological stages. This delay may contribute to the gradual onset of cognitive decline and the development of neuronal pathologies such as neurofibrillary tangles. Specifically, we noted a reduction in the expression of genes associated with cholinergic transmission. The progressive loss of cholinergic afferents in AD is a well-established pathology (Davies and Maloney, 1976), with cholinesterase inhibitors remaining an important treatment option (Dubois et al., 2015; Hampel et al., 2018; Summers et al., 1986). Additionally, recent findings have directly associated the loss of cholinergic afferents with their proximity to plaques (Lee and Chen, 2024). Moreover, the plaque age-associated decrease in synaptic genes is well in line with SILK proteomic experiments in mice showing impaired turnover of synaptic proteins (Hark et al., 2021).

Seemingly, in 10-month-old mice, we managed to capture the initial immune response and increasing immune proliferation from the precipitation of the very first plaques. Surprisingly this increase in immune activity associated genes with increasing plaque was not seen in 18-month-old animals and is therefore likely a rapid response to early deposition (Meyer-Luehmann et al., 2008). Instead, at 18 months, we saw an increase in metabolic-associated genes potentially revealing that plaques demand increased metabolic support as they mature or on the contrary represent mitochondrial dysfunction.

Recent advances in amyloid imaging have introduced structural-specific dyes, facilitating the easy categorization of plaque types through hyperspectral imaging and linear unmixing (Michno et al., 2018; Nystrom et al., 2017; Nystrom et al., 2013; Rasmussen et al., 2017). This method offers an unbiased, more efficient and high-throughput alternative to previous techniques that relied on morphological examination by pathologists. Interestingly, we have identified three distinct populations of amyloid plaques in *App^NL-F^* mice, characterized by their unique structural isoforms. These populations were: Aβ+h+q+, Aβ+h+q-, and Aβ+h-q-, likely corresponding to the morphologically defined dense cored, fibrillar, and diffuse plaques respectively (Dickson and Vickers, 2001). Our findings indicate that in older *App^NL-F^*mice, the majority of plaques exhibit a diffuse and non-pleated structure (Aβ+h-q-/ diffuse populations), which induces lower LAMP1 levels and HOMER1 loss compared to more compact and larger LCO positive plaques. This finding of increased synapse loss with increasing structural maturity is in line with previous findings on morphologically- and thioflavin-defined plaques (Koffie et al., 2009; Masliah et al., 1990; Spires et al., 2005) and also aligns with findings in amyloid positive CU-AP individuals, where diffuse plaques exhibit low levels of neurotoxicity (Dickson et al., 1992; Malek-Ahmadi et al., 2016). Furthermore, the finding of increased neurotoxicity with increasing aggregation is also consistent with studies showing that a higher propensity for Aβ aggregation and the presence of dense cored plaques are associated with increased progression of clinical dementia. (Murray and Dickson, 2014; Rijal Upadhaya et al., 2014; Serrano-Pozo et al., 2016).

The process of precipitating plaque pathology is difficult to capture with conventional, steady state imaging techniques, particularly in comparably late onset mouse models like *App^NL^*^-F^. Dynamic imaging approaches such as chemical or genetic *in vivo* labelling provided important insight, though while in vivo, lack chemical specificity and/or temporal span (Meyer-Luehmann et al., 2008). Using the here employed iSILK-based approach for dynamic imaging of amyloid peptide deposition, our results from 10-month-old (Scheme 1) *App^NL^*^-F^ mice indicate that a plaque center of highly compacted Aβ1-42 forms first (Figure 1 & S1). This is also supported by the investigation of structural plaque isoforms over chronological age showing ∼90% of initial depositions (9-month-old *App^NL^*^-F^) in the hippocampus are already fibrilized (Figure S4). We have seen similar results in a previous iSILK study on the *App^NL-G-F^* model suggesting that plaques initially precipitate as small dense deposits (Michno et al., 2021). However, it seems plaques continue to fibrilize as we see a positive correlation between q-hFTAA binding and increasing plaque age that could represent a dynamic maturation process whereby fibrillar Aβ (hFTAA+) species progress to a more mature state at the center of a plaque. This is of interest as a chronological study on plaque maturation in *App^NL-F^* showed no difference in q/h emission across age though a large variety of plaque maturation stages was observed across plaques within an age group (Parvin et al., 2024).

Delineating the plaque age irrespective of chronological age however facilitates the identification of plaque age specific maturation reported here, highlighting the potential of the dynamic imaging method described here. Surprisingly, however, we also note a large increase in the presence of non-pleated (LCO-negative) plaques with chronological age. Here, at 18 months, ∼80% of plaques (by number) were LCO-negative suggesting that these latterly formed plaques do not deposit through initial core formation. This discrepancy could be explained by the mechanism of plaque compaction and seeding potentially driven by microglial activity (Baik et al., 2016; Huang et al., 2021; Venegas et al., 2017; Yuan et al., 2016). For instance, in older mice, the exceptionally high concentrations of Aβ42 could saturate the normal amyloid seeding and compaction processes. This hypothesis is further supported by studies indicating that in the absence of microglia, amyloid structures tend to be less compact (Kiani Shabestari et al., 2022). A limitation of the current study is that the small, LCO-negative plaques were not captured across two consecutive sections of the iSILK guides transcriptomics experiments, posing a central challenge that should be at the center of future investigations.

Together we present a conceptual and technical innovative study using a novel spatial biology paradigm to detail the earliest events of plaque deposition through plaque maturation. The spatial and temporal resolution along with chemical precision achieved with this method, particularly with respect to the age of the AD mice studied, exceeds steady state technologies and consequently provides a dynamic dimension that would otherwise not be discernible.

## Supporting information

10 Months Enrichment Analysis

18 Months Enrichment Analysis

## Acknowledgements

JH is supported by the NIH (R01 AG078796, R21AG078538, R21AG080705), the Swedish Research Council VR (#2019-02397, #2023-02796), the Swedish Alzheimer Foundation (#AF-968238, #AF-939767, #994281), Race against Dementia/Rosetree Foundation Team Award, the Swedish Brain Foundation (FO2022-0311), Åhlén-Stiftelsen. (#233011), Magnus Bergvalls Stiftelse, Stiftelsen Gamla Tjänarinnor and Gun och Bertil Stohnes Stiftelse. FAE is supported by the Cure Alzheimer Foundation, Alzheimer’s Research UK and the NIH (R01 AG078796). JNS is supported by the (NIH R01 AG078796,R21AG080705). JIW is funded by the Swedish Alzheimer’s Foundation. HZ is a Wallenberg Scholar and a Distinguished Professor at the Swedish Research Council supported by grants from the Swedish Research Council (#2023-00356; #2022-01018 and #2019-02397), the European Union’s Horizon Europe research and innovation program under grant agreement No 101053962, Swedish State Support for Clinical Research (#ALFGBG-71320), the Alzheimer Drug Discovery Foundation (ADDF), USA (#201809-2016862), the AD Strategic Fund and the Alzheimer’s Association (#ADSF-21-831376-C, #ADSF-21-831381-C, #ADSF-21-831377-C, and #ADSF-24-1284328-C), the European Partnership on Metrology, co-financed from the European Union’s Horizon Europe Research and Innovation Programme and by the Participating States (NEuroBioStand, #22HLT07), the Bluefield Project, Cure Alzheimer’s Fund, the Olav Thon Foundation, the Erling-Persson Family Foundation, Familjen Rönströms Stiftelse, Stiftelsen för Gamla Tjänarinnor, Hjärnfonden, Sweden (#FO2022-0270), the European Union’s Horizon 2020 research and innovation programme under the Marie Skłodowska-Curie grant agreement No 860197 (MIRIADE), the European Union Joint Programme – Neurodegenerative Disease Research (JPND2021-00694), the National Institute for Health and Care Research University College London Hospitals Biomedical Research Centre, and the UK Dementia Research Institute at UCL (UKDRI-1003). KB is supported by the Swedish Research Council (#2017–00915 and #2022–00732), the Swedish Alzheimer Foundation (#AF-930351, #AF-939721, #AF-968270, and #AF-994551), Hjärnfonden, Sweden (#FO2017–0243 and #ALZ2022–0006), the Swedish state under the agreement between the Swedish government and the County Councils, the ALF-agreement (#ALFGBG-715986 and #ALFGBG-965240), the European Union Joint Program for Neurodegenerative Disorders (JPND2019–466–236), the Alzheimer’s Association 2021 Zenith Award (ZEN-21–848495), the Alzheimer’s Association 2022–2025 Grant (SG-23– 1038904 QC), La Fondation Recherche Alzheimer (FRA), Paris, France, and the Kirsten and Freddy Johansen Foundation, Copenhagen, Denmark, and Familjen Rönströms Stiftelse, Stockholm, Sweden.

## Author contributions

Conceptualization: JH, JIW, FAE

Methodology: JIW, KS, JG, SK, AS, SD, JH

Investigation: JIW, MD, KS, JG, SK, AS, HBH, JH

Visualization: JIW, MD, KS, JG, SK

Funding acquisition: JH, FAE, JNS

Supervision: JH, FAE, DMC

Writing – original draft: JIW, JH

Writing – review & editing: JIW, MD, KB, HZ, DMC, JNS, FAE, JH

## References

Almstedt, K., Nystrom, S., Nilsson, K.P., and Hammarstrom, P. (2009). Amyloid fibrils of human prion protein are spun and woven from morphologically disordered aggregates. Prion 3, 224–235.

Baik, S.H., Kang, S., Son, S.M., and Mook-Jung, I. (2016). Microglia contributes to plaque growth by cell death due to uptake of amyloid beta in the brain of Alzheimer’s disease mouse model. Glia 64, 2274–2290.

Benitez, D.P., Jiang, S., Wood, J., Wang, R., Hall, C.M., Peerboom, C., Wong, N., Stringer, K.M., Vitanova, K.S., Smith, V.C., et al. (2021). Knock-in models related to Alzheimer’s disease: synaptic transmission, plaques and the role of microglia. Mol Neurodegener 16, 47.

Busche, M.A., and Hyman, B.T. (2020). Synergy between amyloid-beta and tau in Alzheimer’s disease. Nat Neurosci 23, 1183–1193.

Cairns, N.J., Ikonomovic, M.D., Benzinger, T., Storandt, M., Fagan, A.M., Shah, A.R., Reinwald, L.T., Carter, D., Felton, A., Holtzman, D.M., et al. (2009). Absence of Pittsburgh compound B detection of cerebral amyloid beta in a patient with clinical, cognitive, and cerebrospinal fluid markers of Alzheimer disease: a case report. Arch Neurol 66, 1557–1562.

Chen, W.T., Lu, A., Craessaerts, K., Pavie, B., Sala Frigerio, C., Corthout, N., Qian, X., Lalakova, J., Kuhnemund, M., Voytyuk, I., et al. (2020). Spatial Transcriptomics and In Situ Sequencing to Study Alzheimer’s Disease. Cell 182, 976–991 e919.

Choi, H., Lee, E.J., Shin, J.S., Kim, H., Bae, S., Choi, Y., and Lee, D.S. (2023). Spatiotemporal characterization of glial cell activation in an Alzheimer’s disease model by spatially resolved transcriptomics. Exp Mol Med 55, 2564–2575.

Cirrito, J.R., Kang, J.E., Lee, J., Stewart, F.R., Verges, D.K., Silverio, L.M., Bu, G., Mennerick, S., and Holtzman, D.M. (2008). Endocytosis is required for synaptic activity-dependent release of amyloid-beta in vivo. Neuron 58, 42–51.

Davies, P., and Maloney, A.J. (1976). Selective loss of central cholinergic neurons in Alzheimer’s disease. Lancet 2, 1403.

De Strooper, B., and Karran, E. (2016). The Cellular Phase of Alzheimer’s Disease. Cell 164, 603–615.

Dickson, D.W., Crystal, H.A., Mattiace, L.A., Masur, D.M., Blau, A.D., Davies, P., Yen, S.H., and Aronson, M.K. (1992). Identification of normal and pathological aging in prospectively studied nondemented elderly humans. Neurobiol Aging 13, 179–189.

Dickson, T.C., and Vickers, J.C. (2001). The morphological phenotype of beta-amyloid plaques and associated neuritic changes in Alzheimer’s disease. Neuroscience 105, 99–107.

Dubois, B., Chupin, M., Hampel, H., Lista, S., Cavedo, E., Croisile, B., Louis Tisserand, G., Touchon, J., Bonafe, A., Ousset, P.J., et al. (2015). Donepezil decreases annual rate of hippocampal atrophy in suspected prodromal Alzheimer’s disease. Alzheimers Dement 11, 1041–1049.

Ghosal, K., Vogt, D.L., Liang, M., Shen, Y., Lamb, B.T., and Pimplikar, S.W. (2009). Alzheimer’s disease-like pathological features in transgenic mice expressing the APP intracellular domain. P Natl Acad Sci USA 106, 18367–18372.

Habib, N., McCabe, C., Medina, S., Varshavsky, M., Kitsberg, D., Dvir-Szternfeld, R., Green, G., Dionne, D., Nguyen, L., Marshall, J.L., et al. (2020). Disease-associated astrocytes in Alzheimer’s disease and aging. Nat Neurosci 23, 701–706.

Hampel, H., Mesulam, M.M., Cuello, A.C., Farlow, M.R., Giacobini, E., Grossberg, G.T., Khachaturian, A.S., Vergallo, A., Cavedo, E., Snyder, P.J., et al. (2018). The cholinergic system in the pathophysiology and treatment of Alzheimer’s disease. Brain 141, 1917–1933.

Hark, T.J., Rao, N.R., Castillon, C., Basta, T., Smukowski, S., Bao, H., Upadhyay, A., Bomba-Warczak, E., Nomura, T., O’Toole, E.T., et al. (2021). Pulse-Chase Proteomics of the App Knockin Mouse Models of Alzheimer’s Disease Reveals that Synaptic Dysfunction Originates in Presynaptic Terminals. Cell Syst 12, 141–158 e149.

Huang, Y., Happonen, K.E., Burrola, P.G., O’Connor, C., Hah, N., Huang, L., Nimmerjahn, A., and Lemke, G. (2021). Microglia use TAM receptors to detect and engulf amyloid β plaques. Nature immunology 22, 586–594.

Ikonomovic, M.D., Abrahamson, E.E., Price, J.C., Hamilton, R.L., Mathis, C.A., Paljug, W.R., Debnath, M.L., Cohen, A.D., Mizukami, K., DeKosky, S.T., et al. (2012). Early AD pathology in a [C-11]PiB-negative case: a PiB-amyloid imaging, biochemical, and immunohistochemical study. Acta Neuropathol 123, 433–447.

Ikonomovic, M.D., Klunk, W.E., Abrahamson, E.E., Mathis, C.A., Price, J.C., Tsopelas, N.D., Lopresti, B.J., Ziolko, S., Bi, W., Paljug, W.R., et al. (2008). Post-mortem correlates of in vivo PiB-PET amyloid imaging in a typical case of Alzheimer’s disease. Brain 131, 1630–1645.

Kiani Shabestari, S., Morabito, S., Danhash, E.P., McQuade, A., Sanchez, J.R., Miyoshi, E., Chadarevian, J.P., Claes, C., Coburn, M.A., Hasselmann, J., et al. (2022). Absence of microglia promotes diverse pathologies and early lethality in Alzheimer’s disease mice. Cell Rep 39, 110961.

Koffie, R.M., Meyer-Luehmann, M., Hashimoto, T., Adams, K.W., Mielke, M.L., Garcia-Alloza, M., Micheva, K.D., Smith, S.J., Kim, M.L., Lee, V.M., et al. (2009). Oligomeric amyloid beta associates with postsynaptic densities and correlates with excitatory synapse loss near senile plaques. P Natl Acad Sci USA 106, 4012–4017.

Koutarapu, S., Ge, J., Dulewicz, M., Srikrishna, M., Szadziewska, A., Wood, J., Blennow, K., Zetterberg, H., Michno, W., Ryan, N.S., et al. (2024). Chemical signatures delineate heterogeneous amyloid plaque populations across the Alzheimer’s disease spectrum. bioRxiv.

Lee, M.K., and Chen, G. (2024). Loss of Cholinergic and Monoaminergic Afferents in APPswe/PS1DeltaE9 Transgenic Mouse Model of Cerebral Amyloidosis Preferentially Occurs Near Amyloid Plaques. Int J Mol Sci 25.

Long, J.M., and Holtzman, D.M. (2019). Alzheimer Disease: An Update on Pathobiology and Treatment Strategies. Cell 179, 312–339.

Malek-Ahmadi, M., Perez, S.E., Chen, K., and Mufson, E.J. (2016). Neuritic and Diffuse Plaque Associations with Memory in Non-Cognitively Impaired Elderly. J Alzheimers Dis 53, 1641–1652.

Mallach, A., Zielonka, M., van Lieshout, V., An, Y., Khoo, J.H., Vanheusden, M., Chen, W.T., Moechars, D., Arancibia-Carcamo, I.L., Fiers, M., et al. (2024). Microglia-astrocyte crosstalk in the amyloid plaque niche of an Alzheimer’s disease mouse model, as revealed by spatial transcriptomics. Cell Rep 43, 114216.

Masliah, E., Terry, R.D., Mallory, M., Alford, M., and Hansen, L.A. (1990). Diffuse plaques do not accentuate synapse loss in Alzheimer’s disease. The American journal of pathology 137, 1293–1297.

Matarin, M., Salih, D.A., Yasvoina, M., Cummings, D.M., Guelfi, S., Liu, W., Nahaboo Solim, M.A., Moens, T.G., Paublete, R.M., Ali, S.S., et al. (2015). A genome-wide gene-expression analysis and database in transgenic mice during development of amyloid or tau pathology. Cell Rep 10, 633–644.

Meyer-Luehmann, M., Spires-Jones, T.L., Prada, C., Garcia-Alloza, M., de Calignon, A., Rozkalne, A., Koenigsknecht-Talboo, J., Holtzman, D.M., Bacskai, B.J., and Hyman, B.T. (2008). Rapid appearance and local toxicity of amyloid-beta plaques in a mouse model of Alzheimer’s disease. Nature 451, 720–724.

Michno, W., Kaya, I., Nystrom, S., Guerard, L., Nilsson, K.P.R., Hammarstrom, P., Blennow, K., Zetterberg, H., and Hanrieder, J. (2018). Multimodal Chemical Imaging of Amyloid Plaque Polymorphism Reveals Abeta Aggregation Dependent Anionic Lipid Accumulations and Metabolism. Anal Chem 90, 8130–8138.

Michno, W., Stringer, K.M., Enzlein, T., Passarelli, M.K., Escrig, S., Vitanova, K., Wood, J., Blennow, K., Zetterberg, H., Meibom, A., et al. (2021). Following spatial Abeta aggregation dynamics in evolving Alzheimer’s disease pathology by imaging stable isotope labeling kinetics. Sci Adv 7, eabg4855.

Morris, G.P., Clark, I.A., and Vissel, B. (2014). Inconsistencies and controversies surrounding the amyloid hypothesis of Alzheimer’s disease. Acta Neuropathol Commun 2, 135.

Murray, M.E., and Dickson, D.W. (2014). Is pathological aging a successful resistance against amyloid-beta or preclinical Alzheimer’s disease? Alzheimers Res Ther 6, 24.

Nilsson, K.P., Aslund, A., Berg, I., Nystrom, S., Konradsson, P., Herland, A., Inganas, O., Stabo-Eeg, F., Lindgren, M., Westermark, G.T., et al. (2007). Imaging distinct conformational states of amyloid-beta fibrils in Alzheimer’s disease using novel luminescent probes. Acs Chem Biol 2, 553–560.

Nystrom, S., Back, M., Nilsson, K.P.R., and Hammarstrom, P. (2017). Imaging Amyloid Tissues Stained with Luminescent Conjugated Oligothiophenes by Hyperspectral Confocal Microscopy and Fluorescence Lifetime Imaging. Jove-J Vis Exp.

Nystrom, S., Psonka-Antonczyk, K.M., Ellingsen, P.G., Johansson, L.B., Reitan, N., Handrick, S., Prokop, S., Heppner, F.L., Wegenast-Braun, B.M., Jucker, M., et al. (2013). Evidence for age-dependent in vivo conformational rearrangement within Abeta amyloid deposits. Acs Chem Biol 8, 1128–1133.

Parvin, F., Haglund, S., Wegenast-Braun, B., Jucker, M., Saito, T., Saido, T.C., Nilsson, K.P.R., Nilsson, P., Nystrom, S., and Hammarstrom, P. (2024). Divergent Age-Dependent Conformational Rearrangement within Abeta Amyloid Deposits in APP23, APPPS1, and App(NL-F) Mice. ACS Chem Neurosci 15, 2058–2069.

Rasmussen, J., Mahler, J., Beschorner, N., Kaeser, S.A., Hasler, L.M., Baumann, F., Nystrom, S., Portelius, E., Blennow, K., Lashley, T., et al. (2017). Amyloid polymorphisms constitute distinct clouds of conformational variants in different etiological subtypes of Alzheimer’s disease. P Natl Acad Sci USA 114, 13018–13023.

Rijal Upadhaya, A., Kosterin, I., Kumar, S., von Arnim, C.A., Yamaguchi, H., Fandrich, M., Walter, J., and Thal, D.R. (2014). Biochemical stages of amyloid-beta peptide aggregation and accumulation in the human brain and their association with symptomatic and pathologically preclinical Alzheimer’s disease. Brain 137, 887–903.

Rohr, D., Boon, B.D.C., Schuler, M., Kremer, K., Hoozemans, J.J.M., Bouwman, F.H., El-Mashtoly, S.F., Nabers, A., Grosserueschkamp, F., Rozemuller, A.J.M., et al. (2020). Label-free vibrational imaging of different Abeta plaque types in Alzheimer’s disease reveals sequential events in plaque development. Acta Neuropathol Commun 8, 222.

Saito, T., Matsuba, Y., Mihira, N., Takano, J., Nilsson, P., Itohara, S., Iwata, N., and Saido, T.C. (2014). Single App knock-in mouse models of Alzheimer’s disease. Nature neuroscience 17, 661–663.

Sasaguri, H., Nilsson, P., Hashimoto, S., Nagata, K., Saito, T., De Strooper, B., Hardy, J., Vassar, R., Winblad, B., and Saido, T.C. (2017). APP mouse models for Alzheimer’s disease preclinical studies. The EMBO journal 36, 2473–2487.

Serrano-Pozo, A., Betensky, R.A., Frosch, M.P., and Hyman, B.T. (2016). Plaque-Associated Local Toxicity Increases over the Clinical Course of Alzheimer Disease. The American journal of pathology 186, 375–384.

Sims, J.R., Zimmer, J.A., Evans, C.D., Lu, M., Ardayfio, P., Sparks, J., Wessels, A.M., Shcherbinin, S., Wang, H., Monkul Nery, E.S., et al. (2023). Donanemab in Early Symptomatic Alzheimer Disease: The TRAILBLAZER-ALZ 2 Randomized Clinical Trial. JAMA 330, 512–527.

Smith, K.D., Prince, D.K., MacDonald, J.W., Bammler, T.K., and Akilesh, S. (2024). Challenges and Opportunities for the Clinical Translation of Spatial Transcriptomics Technologies. Glomerular Dis 4, 49–63.

Spires, T.L., Meyer-Luehmann, M., Stern, E.A., McLean, P.J., Skoch, J., Nguyen, P.T., Bacskai, B.J., and Hyman, B.T. (2005). Dendritic spine abnormalities in amyloid precursor protein transgenic mice demonstrated by gene transfer and intravital multiphoton microscopy. J Neurosci 25, 7278–7287.

Summers, W.K., Majovski, L.V., Marsh, G.M., Tachiki, K., and Kling, A. (1986). Oral tetrahydroaminoacridine in long-term treatment of senile dementia, Alzheimer type. N Engl J Med 315, 1241–1245.

van Dyck, C.H., Swanson, C.J., Aisen, P., Bateman, R.J., Chen, C., Gee, M., Kanekiyo, M., Li, D., Reyderman, L., Cohen, S., et al. (2023). Lecanemab in Early Alzheimer’s Disease. N Engl J Med 388, 9–21.

Venegas, C., Kumar, S., Franklin, B.S., Dierkes, T., Brinkschulte, R., Tejera, D., Vieira-Saecker, A., Schwartz, S., Santarelli, F., Kummer, M.P., et al. (2017). Microglia-derived ASC specks cross-seed amyloid-beta in Alzheimer’s disease. Nature 552, 355–361.

Williams, J.B., Cao, Q., and Yan, Z. (2021). Transcriptomic analysis of human brains with Alzheimer’s disease reveals the altered expression of synaptic genes linked to cognitive deficits. Brain Commun 3, fcab123.

Wood, J.I., Wong, E., Joghee, R., Balbaa, A., Vitanova, K.S., Stringer, K.M., Vanshoiack, A., Phelan, S.J., Launchbury, F., Desai, S., et al. (2022). Plaque contact and unimpaired Trem2 is required for the microglial response to amyloid pathology. Cell Rep 41, 111686.

Yuan, P., Condello, C., Keene, C.D., Wang, Y., Bird, T.D., Paul, S.M., Luo, W., Colonna, M., Baddeley, D., and Grutzendler, J. (2016). TREM2 Haplodeficiency in Mice and Humans Impairs the Microglia Barrier Function Leading to Decreased Amyloid Compaction and Severe Axonal Dystrophy. Neuron 90, 724–739.

