## Supplementary Figures for "Isotope Encoded chemical Imaging Identifies Amyloid Plaque Age Dependent Structural Maturation, Synaptic Loss, and Increased Toxicity"

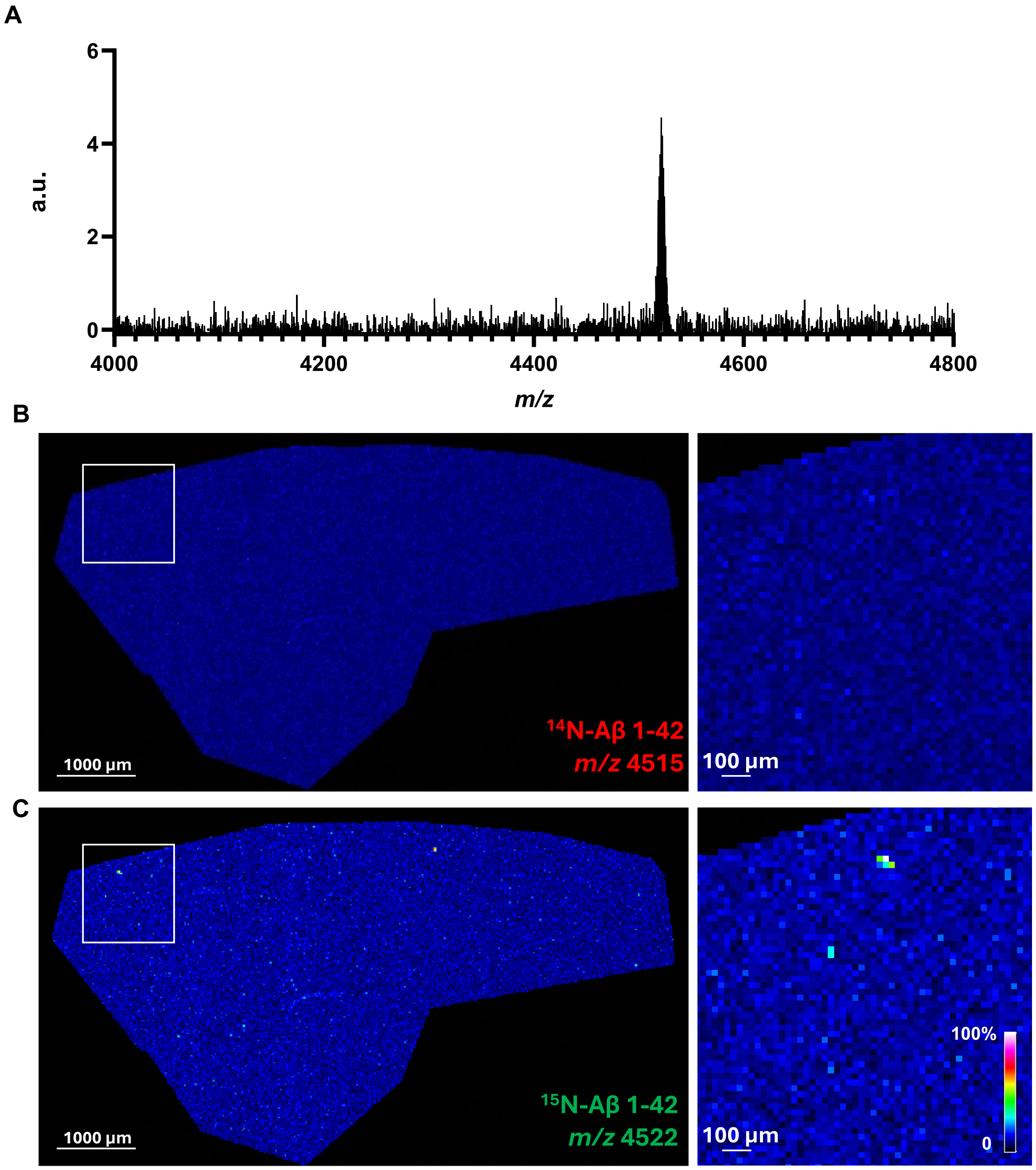


Figure S1. MALDI MSI analysis revealed that plaques in 10-month *App*$^{NL\text{-}F}$ mice contained only $^{15}$N-labeled Aβ 1-42, with no unlabelled Aβ 1-42 detected. (A) The spectrum from MALDI MSI showed that the plaques in 10-month *App*$^{NL\text{-}F}$ mice contained solely Aβ 1-42. (B, C) Single ion images of Aβ 1-42 (B) showing no unlabelled ($^{14}$N) Aβ 1-42 (C) only $^{15}$N-labeled Aβ 1-42 was detected.

10 months old included for analysis

| N | AOI | Peak Start | Peak End | Centroid | Local Intersection |
|---|---|---|---|---|---|
| 1 | 1 | 4515,19 | 4535,04 | 4524,92 | 9,79 |
| 2 | 2 | 4515,75 | 4535,23 | 4525,42 | 9,64 |
| 3 | 3 | 4514,81 | 4534,67 | 4525,16 | 10,09 |
| 4 | 5 | 4512,94 | 4535,60 | 4524,28 | 11,24 |
| 5 | 6 | 4515,94 | 4532,42 | 4525,15 | 8,46 |
| 6 | 9 | 4516,82 | 4537,32 | 4525,84 | 9,70 |
| 7 | 10 | 4514,14 | 4538,27 | 4526,12 | 12,07 |
| 8 | 12 | 4517,21 | 4537,51 | 4526,17 | 9,46 |
| 9 | 13 | 4515,10 | 4535,78 | 4527,39 | 11,31 |
| 10 | 17 | 4508,12 | 4529,00 | 4519,31 | 10,87 |
| 11 | 18 | 4510,04 | 4528,81 | 4519,01 | 9,10 |
| 12 | 19 | 4505,83 | 4530,72 | 4519,2 | 13,18 |
| 13 | 20 | 4511,38 | 4530,72 | 4519,39 | 8,72 |
| 14 | 21 | 4510,81 | 4529,38 | 4519,8 | 9,08 |
| 15 | 22 | 4506,78 | 4529,38 | 4519,07 | 11,97 |
| 16 | 23 | 4510,04 | 4531,30 | 4520,5 | 10,51 |

18 months old included for analysis

| N | AOI | Peak Start | Peak End | Centroid | Local Intersection |
|---|---|---|---|---|---|
| 1 | 1 | 4507,19 | 4526,92 | 4517,12 | 9,93 |
| 2 | 2 | 4506,62 | 4525,77 | 4516,18 | 9,56 |
| 3 | 3 | 4506,24 | 4527,49 | 4517,08 | 10,85 |
| 4 | 5 | 4505,66 | 4526,92 | 4517,01 | 11,35 |
| 5 | 6 | 4507,00 | 4524,81 | 4515,64 | 8,64 |
| 6 | 7 | 4510,32 | 4526,99 | 4518,47 | 8,15 |
| 7 | 8 | 4510,13 | 4529,24 | 4519,53 | 9,40 |
| 8 | 9 | 4507,51 | 4529,05 | 4518,95 | 11,44 |
| 9 | 10 | 4510,13 | 4525,49 | 4518,20 | 8,07 |
| 10 | 11 | 4508,45 | 4528,67 | 4518,99 | 10,54 |
| 11 | 12 | 4512,75 | 4527,74 | 4519,78 | 7,03 |
| 12 | 14 | 4508,70 | 4528,05 | 4518,46 | 9,76 |
| 13 | 16 | 4511,00 | 4526,32 | 4518,21 | 7,21 |
| 14 | 17 | 4510,24 | 4525,94 | 4517,97 | 7,73 |
| 15 | 19 | 4504,05 | 4524,93 | 4514,83 | 10,78 |
| 16 | 20 | 4505,01 | 4524,93 | 4515,03 | 10,02 |
| 17 | 21 | 4504,82 | 4525,88 | 4515,18 | 10,36 |
| 18 | 24 | 4507,11 | 4526,27 | 4515,95 | 8,83 |

**Table S1.** MALDI MSI spectral data for Aβ1-42 signals across single plaques in 10- and 18-month-old $App^{NL\text{-}F}$ mice.

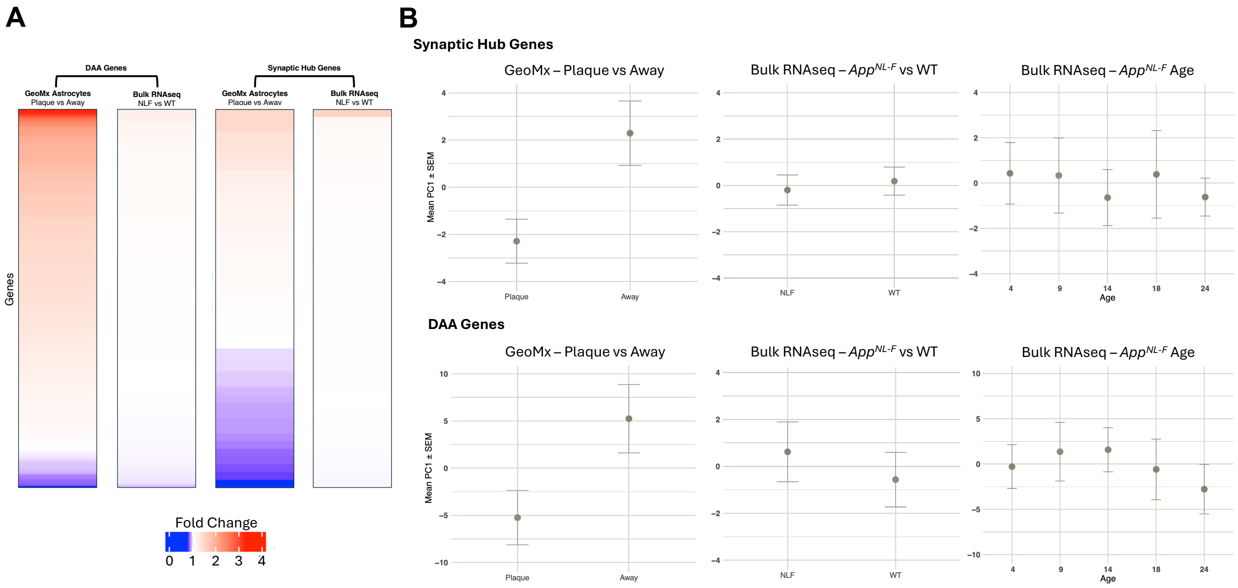


Figure S2. Amyloid induced changes in gene expression detected using either GeoMx spatial transcriptomics or Bulk RNAseq. (A) Heatmaps comparing fold change of Disease Associated Astrocytic genes (DAAs, Habib et al., 2020) and synaptic hub genes (Williams et al., 2021) by use of GeoMx spatial transcriptomic technology using an astrocytic collection in plaque vs non-plaque associated areas (n=6) or by use of bulk hippocampal RNA sequencing in 18-month-old wild-type (n=11) vs *App*$^{NL\text{-}F}$ (n=9) mice. (B) Principal Component Analysis of the DAAs and synaptic hub genes performed using data from both GeoMx spatial transcriptomics and Bulk hippocampal RNA sequencing.

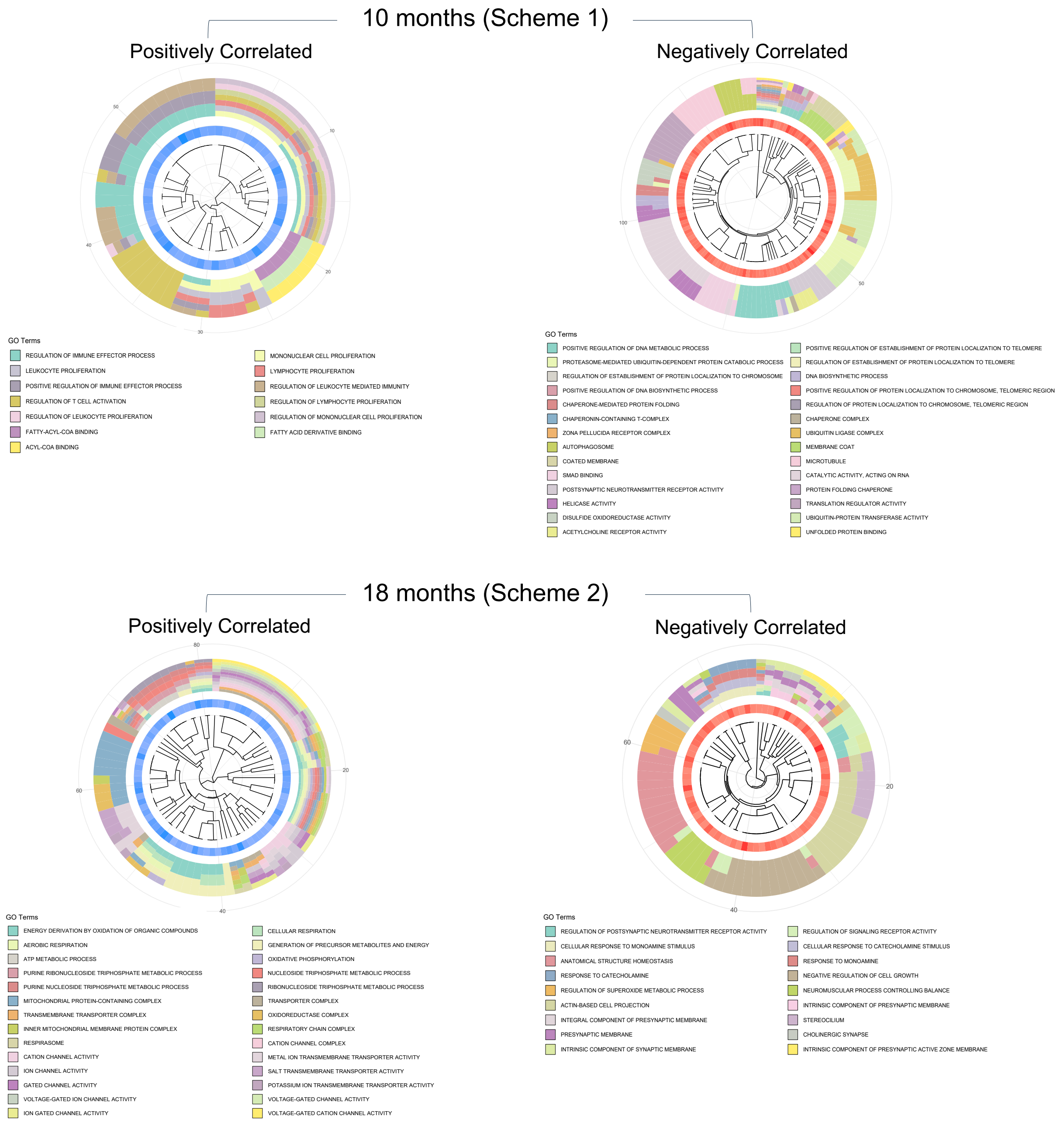


Figure S3. Gene Ontology Cluster plots for genes correlated with plaque age in 10-month-old (Scheme 1) and 18-month-old (Scheme 2) *App*$^{NL\text{-}F}$ mice.

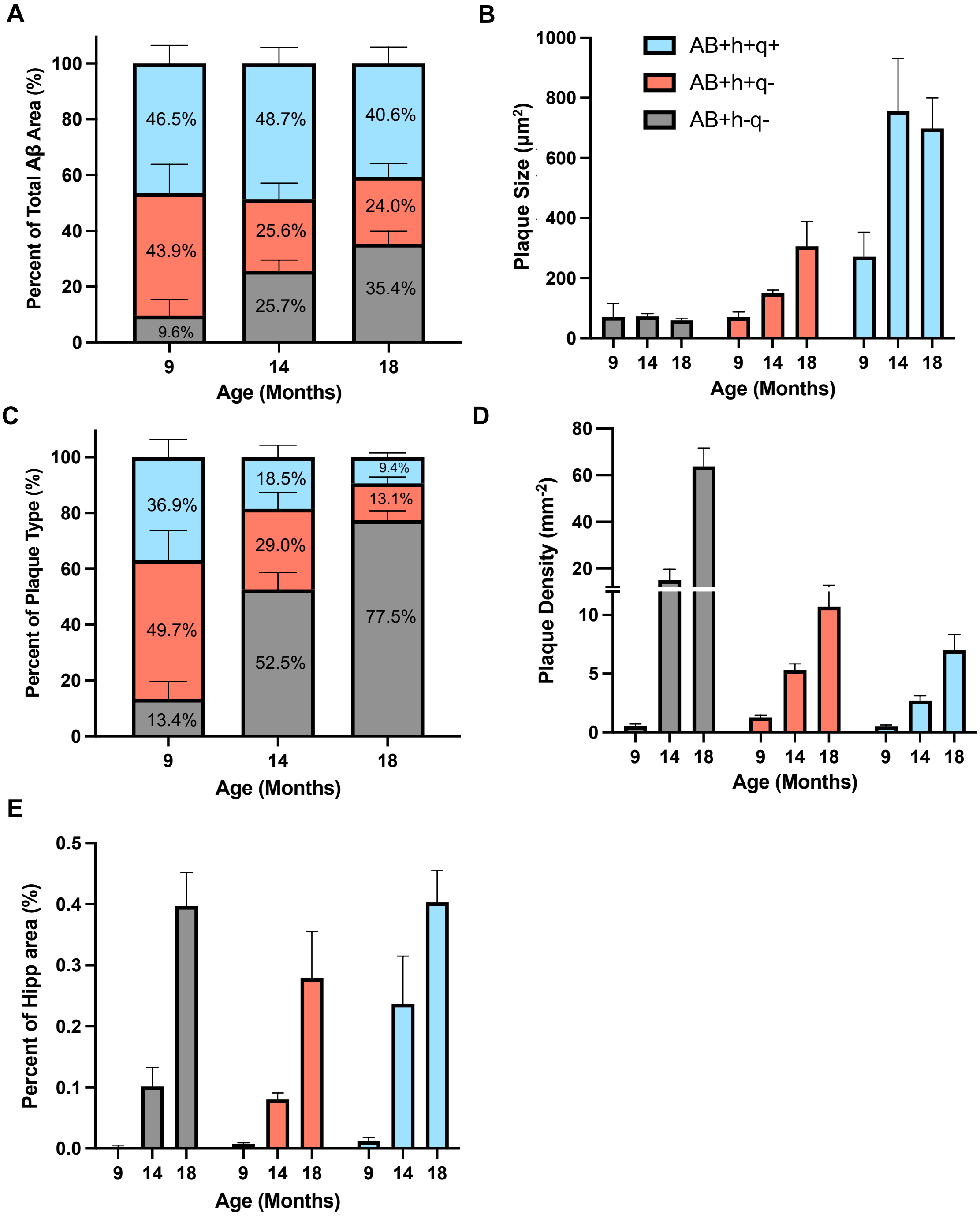


Figure S4. Plaque type characterization in 9-, 14-, and 18-month-old *App*$^{NL-F}$ mice (A) Distribution of plaque types as a percentage of the total Aβ positive area. (B) Average area occupied by each plaque type. (C) Distribution of plaque types as a percentage of the total number of Aβ plaques. (D) Plaque density of each plaque type, measured as the number of plaques per unit area in the hippocampus. (E) Percentage of the hippocampal area occupied by each plaque type. 18 months n=6 (data also shown in Figure 4), 14 months n=5, 9 months n=7 (a total of n=14 were tested, only n=7 were plaque positive in the hippocampus).
